# Post-mortem evidence for a reciprocal relationship between genomic DNA damage and alpha-synuclein pathology in dementia with Lewy bodies

**DOI:** 10.1101/2024.04.24.590825

**Authors:** David J. Koss, Olivia Todd, Hariharan Menon, Zoe Anderson, Tamsin Yang, Johannes Attems, Fiona E. LeBeau, Daniel Erskine, Tiago F. Outeiro

## Abstract

DNA damage and DNA damage repair (DDR) dysfunction are insults with broad implications on cellular physiology, including in proteostasis, and have been recently implicated in many neurodegenerative diseases. Alpha-synuclein (aSyn), a pre-synaptic and nuclear protein associated with neurodegenerative disorders known as synucleinopathies, has been implicated in DNA double strand break (DSB) repair function. Consistently, DSB induction has been demonstrated in cell and animal models of synucleinopathy. Nevertheless, the types of DNA damage and the contribution of DNA damage towards Lewy body (LB) formation in synucleinopathies are unknown. Here, we demonstrate the increase of DSB in neuronal and non-neuronal cellular populations of post-mortem temporal cortex tissue from dementia with Lewy body (DLB) patients and demonstrate increases in DSBs early at a presymptomatic age of aSyn transgenic mice. Strikingly, in postmortem DLB tissue, DNA damage-derived ectopic cytoplasmic genomic material (eCGM) was evident within the majority of LBs examined. The observed cellular pathology was consistent with nucleoproteasomal upregulation of associated DNA damage repair proteins, particularly in base excision repair and DSB repair pathways. Collectively our study demonstrates the early occurrence of DNA damage and associated nucleoproteasomal changes in response to nuclear aSyn pathology. Furthermore, the data suggests a potential involvement for DNA damage derived eCGM for the facilitation of cytoplasmic aSyn aggregates. Ultimately, uncovering pathological mechanisms underlying DNA damage in DLB sheds light into novel disease mechanisms and opens novel possibilities for diagnosing and treating synucleinopathies.

## Introduction

The maintenance of accurate genetic information for the process of transcription and protein translation is essential for cellular homeostasis, particularly in post-mitotic, long lived central nervous system (CNS) neurons. Nevertheless, numerous DNA-damaging insults occur frequently, possibly leading to a myriad of disruptive genomic alterations, including base modifications, single strand (SSBs) and double strand breaks (DSBs), all of which can interrupt transcription [13, 28]. Thus, maintaining genomic integrity requires rapid engagement of DNA damage repair (DDR) pathways to deal with insults. These DDR pathways include base modification and SSBs repair, via base-excision repair (BER), and DSB repair, via non-homologous end-joining (NHEJ) and homologous recombination (HR) [10]. Owing to the restriction of high-fidelity HR repair to specific cell cycle stages, the capacity of neurons to maintain genomic integrity is limited to the more error-prone NHEJ for DSB repair pathways [42]. Consequently, aging neurons exhibit an increased number of individual genomic errors, leading to the accumulation of somatic mutations within the brain, compromising optimal neuronal functioning and increasing the risk for disease [33, 37].

In addition, excessive unresolved DNA damage has been linked to senescence and the activation of canonical cell death pathways including apoptosis, but also the DNA damage specific pathway Parthantosis [61]. Equally, DNA damage can result in the generation of extra-nuclear genomic material in the form of micronuclei and chromatin cytoplasmic fragments[44]. Such ectopic cytoplasmic genomic material (eCGM) can aggravate inflammatory processes[44] but also may lead to aberrant protein-chromatin interactions, with pro-aggregation properties for many neurodegenerative relevant proteins [11, 22, 71].

Given that DNA damage can be considered one of the primary biological substrates of age [81], and the vulnerability of CNS neurons to accumulate such damage, it is unsurprising that DNA damage and genomic instability are now emerging as cellular stressors central to the etiopathology of many age-related neurodegenerative diseases, including Alzheimer’s disease (AD), amyotrophic lateral sclerosis (ALS), and Parkinson’s disease (PD) [19, 57, 62].

In relation to synucleinopathies, a number of studies have reported on the accumulation of base oxidation, SSB and DSB in post-mortem PD tissue [1, 24, 56, 57, 82]. Parkinson’s disease (PD) and dementia with Lewy bodies (DLB) share, as a common pathological hallmark, the accumulation of protein-rich inclusions known as Lewy bodies (LBs) within cortical and brain stem neurons of affected individuals. Alpha-synuclein (aSyn) is the main protein component of LBs, and excessive DNA damage and DDR impairment has been reported in association with aSyn pathology in various model systems, particularly in relation to DSB formation and repair [51, 53, 57, 76, 80]. However, whether DNA damage and alteration in DDR are relevant in DLB has not as yet been established. Importantly, such alterations are compatible with both loss of physiological and gain of toxic aSyn functions. Consistently, both pathological modification of aSyn and *SNCA* gene knockout in rodents have been associated with a reduced capacity for DNA damage repair, leading to elevated levels of DSBs and downstream neurodegeneration [57, 79]. Strikingly, we recently demonstrated the occurrence of nuclear aSyn in postmortem human tissue and the occurrence of disease-dependent modifications of nuclear aSyn in DLB cases [34]. Nevertheless, the consequences of nuclear aSyn pathology upon genomic stability, DNA damage and the ensuing changes in nuclear proteasome have yet to be determined.

Here, employing fixed and frozen post-mortem human tissue of clinical and neuropathologically diagnosed cases of DLB, we investigate multiple aspects of DNA damage and nuclear pathology in relation to this disease. The study provides a quantification of DNA damage within neuronal and non-neuronal cortical populations, evidence for the involvement of eCGM in LB formation and a characterisation of nuclear proteasomal changes in DLB. Collectively the work further highlights the involvement of DNA damage and DDR in the etiopathology of synucleinopathies.

## Materials and Methods

### Human post-mortem brain tissue

Brain tissue from clinical and neuropathologically confirmed human cases of DLB and non-neurodegenerative disease controls (Con) was obtained from the Newcastle Brain tissue resource (NBTR). Slide mounted fixed paraffin embedded tissue sections from the middle and superior temporal gyrus (BA21/22) were used for immunohistochemistry (Con, n=13 and DLB, n=12). Frozen temporal cortex (BA21/22) from the contralateral hemisphere was used for subcellular fractionation and subsequent western blots (n=12 for both Con and DLB) and for mass spectrometry (n=9 for both Con and DLB).

Disease confirmation for each case was achieved by a review of clinical history upon death and detailed neuropathological assessment according to Braak Lewy body stages [5], the McKeith Criteria [40, 41] and the National institute of ageing -Alzheimer’s association (NIA-AA) criteria [47] inclusive of Thal phases[72], Braak neurofibrillary tangle stages[4] and the Consortium to Establish a Registry for Alzheimer’s Disease (CERAD) scoring for neuritic plaques[46]. Collectively there was no significant difference in either age or post-mortem delay between Con and DLB cases (p>0.05). See table 1 and supplementary table 1 for full details.

**Table 1.**
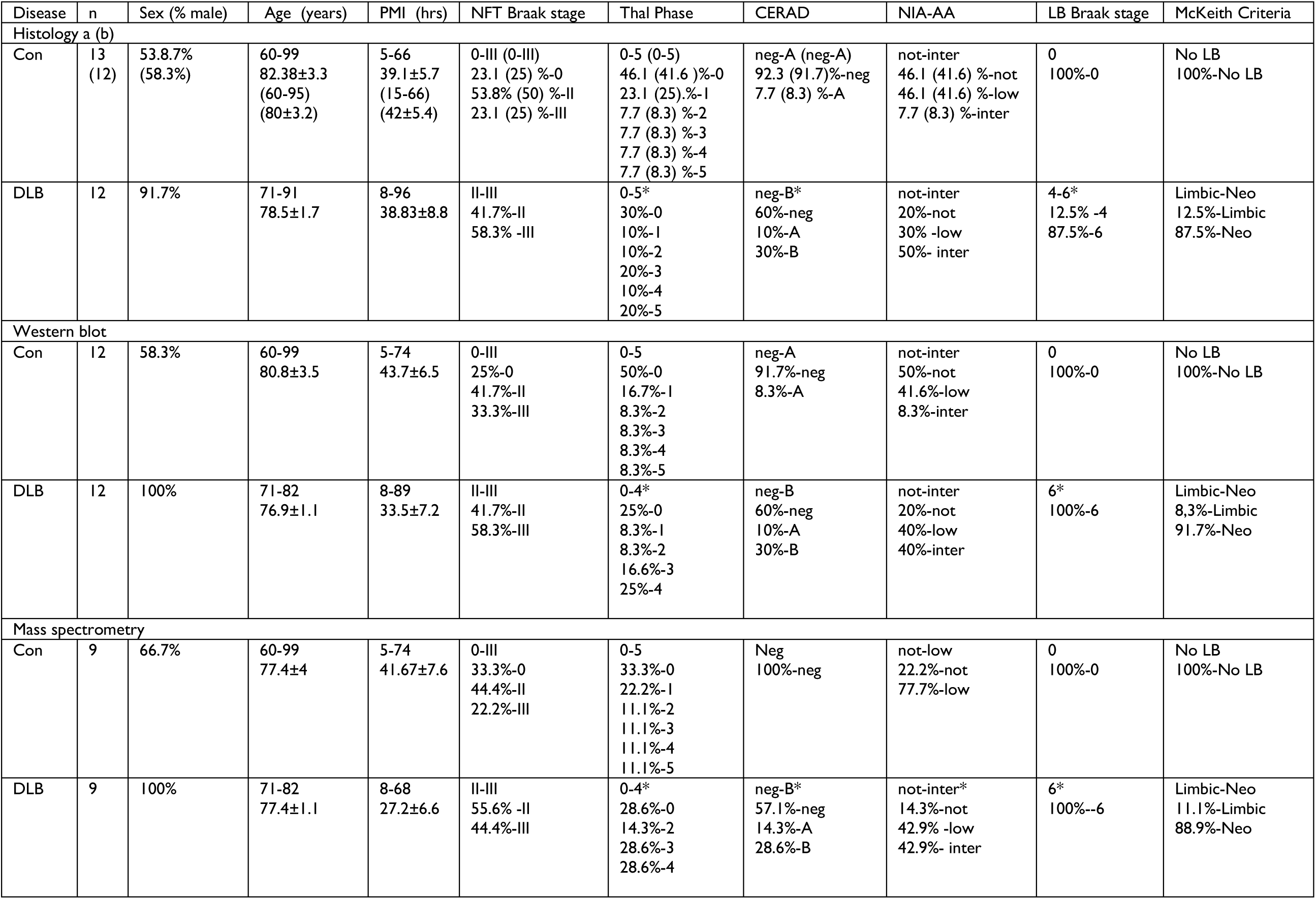
Human tissue cohort. Human cases use for histology, western blot and mass spectrometry. Cases are separated by disease classification according to non-diseased controls (Con) and dementia with Lewy bodies (DLB) per methodological protocol (Histology, Western blot and Mass spectrometry). Case numbers (n), sex, age, post-mortem interval (PMI), neurofibrillary tangle (NFT) Braak stage, Thal phase, Consortium to Establish a Registry for Alzheimer’s Disease (CERAD), the National Institute of Ageing – Alzheimer’s Association (NIA-AA) criteria, Lewy body (LB) Braak stage and McKeith criteria are provided. For age and PMI both range and mean ±SEM are provided. For numerical scores of pathology, range and percentage composition are given. For CERAD scores, negative (neg), A and B reported. For NIA-AA, not, low and intermediate (inter) risk for Alzheimer’s disease. For McKeith criteria, only percentage composition is given, where cases free of LBs (No LB), Limbic and neocortical (Neo) predominant are indicated. a= cohort for immunohistochemical assessment of γH2A.X and XRCC1 and b= cohort for immunohistochemical assessment of γH2A.X and nuclear pS129-aSyn.

### Animal brain tissue

C57BL/6 wild type (WT) and A30P aSyn transgenic mice were used for immunohistochemical quantification of DNA damage. A30P mice express human mutant aSynA30P, under the control of the Thy-1 promotor [30], and were bred in-house from homozygous breeding pairs originally supplied by Dr. P. Kahle (University of Tubingen). Animals were rederived to a C57Bl/6 background via breeding with C57BL/6 mice (Charles River Laboratories, Tranent, UK) and maintained as previously described[73]. C57BL/6 WT control mice were used for comparison and purchased from Charles Rivers laboratories. In-house, as per the ARRIVE guidelines, all mice were group housed (3-4 mice per cage), on a 12 hour light/dark cycle, lights on at 7:00 am, with food and water provided ad libitum.

At 2.5-3 months of age (6-8 weeks of age), male mice were anesthetised with isoflurane via inhalation, prior to intra-muscular injection of Ketamine (> 100 mg/kg) and Xylazine (> 10 mg/kg) and euthanised by transcranial perfusion of 4% paraformaldehyde (PFA; Thermofisher). Fixed whole brains were removed, stored in PFA at 4°C overnight, before being transferred to cryoprotectant (30% glycerol and 30% ethylene glycol in 0.3M PBS in ddH_2_O) at 4°C overnight and finally stored at −20°C, prior to use. All experiments were performed in accordance with the UK animals (Scientific Procedures) Act 1986 and European Union directive 2010/63EU.

### Immunohistochemical staining of human post-mortem brain tissue

Paraffin embedded fixed slide mounted tissue sections (10 µm thick) were baked at 60°C for 30mins, dewaxed in Xylene (2 x 5mins) and rehydrated via submersion in descending concentrations of ethanol (99, 95 and 70%, for 5mins). Slides were washed in Tris-buffered Saline (TBS; 5mM Tris, 145mM NaCl, pH 7.4) for 5 mins, prior to antigen retrieval. Optimised nuclear staining as previously determined [34] was achieved via pressured heating in EDTA (10mM, pH 8) for 2mins, prior to submersion in formic acid (90%, 10mins) at RT. Slides were washed in TBS x2 and in TBS with 0.1 % Tween-20 (TBST) for 5mins, prior to being blocked in TBST containing 10% normal goat serum (NBS, Sigma) for 1hr at RT. Primary antibodies: mouse anti-γH2A.X IgG1 (Clone JBW301, Cat# 05-636, Sigma, 1:500); rabbit anti-XRCC1 (Cat# HPA0061717, Sigma, 1:250) or rabbit anti-pS129-aSyn (EP1536Y, Cat# ab15253, abcam, 1:500) and mouse anti-NeuN IgG2b (Cat# ab104223, abcam, 1:250), were applied to slides in 10 % NGS containing TBST overnight at 4°C. Following 3 x washes in TBST, slides were incubated in secondary antibodies; goat anti-mouse IgG1 Alexa 488 (1:500, Fisher Scientific), goat anti-mouse IgG2b Alexa 647 (1:500, Fisher Scientific) and goat anti-rabbit Alexa 594 (1:500, Fisher Scientific) in 10% NGS containing TBST for 1hr at RT. Following 3x washes in TBST and 30sec incubation in 70% ethanol, autofluorescence was quenched in 0.03% Sudan Black B in 70% ethanol for 5mins. Stained slides were coverslipped with Prolong Diamond Mountant with DAPI (Fisher Scientific) and stored in the dark at −20°C before being imaged.

### Immunohistochemical staining of mouse brain tissue

Fixed brains from WT (n=5) and homozygous A30P (n=5) brains from 2.5-3-month male old mice were removed from cryoprotectant solutions, hemisected and cut into 35 µM thick sections by means of a freezing microtome and stored in 1M PBS at 4°C prior to staining.

Free-floating brain sections were washed three times in 1M PBS and subsequently blocked for five hours submerged in 10% NGS containing 1M PBS with 1% triton (PBS-T), on a vibrating (160 rpm) shaker (Grant – bio PMS-1000i) at RT. Sections were then incubated in 10% NGS PBS-T with primary antibodies for XRCC-1, γH2A.X and NeuN (as above) overnight at 4°C with continued shaking (160 rpm). Following 3 times washes in PBS at RT, sections were incubated with appropriate secondary antibodies (as detailed above) for 3hrs (RT, 160rpm). Autofluorescence was then quenched, incubating sections in 70 % ethanol for 5 mins, prior to submersion in 0.03% Sudan black B in 70% ethanol for 5 mins and washed 3 times in PBS. The last PBS wash contained the nuclear acid Hoechst 33342 dye (2 µg/ml). Sections were then transferred directly to superfrost plus slides and mounted in Fluoromout-G (Thermofisher) and allowed to dry overnight. Stained slide mounted sections were stored in the dark at −20°C, prior to being imaged.

### Nuclear immunofluorescence tissue staining

Immunofluorescence-stained post-mortem human and mouse brain tissue slides were imaged either on a widefield fluorescence (Nikon Eclispse 90i microscope, DsQ1Mc camera and NIS element software V 3.0, Nikon) or confocal (Leica SP 8, LAS Z software, Lecia -microsystems) microscope. Quantifications of XRCC-1 and γH2A.X in human and mouse tissues as well as γH2A.X and pS129-aSyn in human sections were conducted via Leica SP 8 confocal microscope under 63x oil immersion lens. In human slides, 5 fields of interest were selected at random from the lateral temporal grey matter (cortical layers V-VI) and 3 fields of interest were selected at random in mouse cortical sections (Somatosensory cortex). Per region, Z-stacks (21 images at 0.3 µm step thickness totalling a depth of 6.3 µm depth) were captured for XRCC-1, γH2A.X, NeuN immunoreactivity and DAPI or Hoechst fluorescence for each sample (For XRCC-1 and γH2A.X, laser setting in human: 12.1 %, 405nm, 14.2 % 488 nm, 1.4 % 552 nm 3.2% 638 nm and in mouse: 8 % 405 nm; 30.5 % 488 nm; 25 % 552 nm and 2.3% 638 nm and for γH2A.X and pS129-aSyn in human: 7.6 % 405 nm; 8% 488nm; 8.6 % 552 nm and 1.9 % 638 nm).

For imaging, Z-stacks were summated using Z-project maximum intensity function of ImageJ (NIH), DAPI fluorescence was used to manually trace individual nuclei as regions of interest (ROIs), and corresponding intensity of NeuN staining within each ROI used to divide ROIs into NeuN positive (NeuN +ve, neuronal cells) and NeuN negative (NeuN -ve, non-neuronal cells). Mean fluorescence intensity values for γH2A.X, XRCC-1 and pS129 aSyn, without threshold application, were calculated per ROIs and pooled into either NeuN +ve and NeuN -ve generating a mean value per image per immunoreactive channel. Values per image were then further averaged to determine a mean value of fluorescence per channel for NeuN +ve and NeuN -ve nuclei per case. The resulting values were then pooled into disease conditions or genotype prior to comparative analysis. Disease or genotype comparisons (e.g Con Cf. DLB and WT cf. A30P) of channel fluorescence as well as raw counts NeuN +ve and NeuN -ve nuclei number where performed via Mann-Whitney tests (GraphPad Prism, Ver 5). For mouse comparisons, owing to the existence of a clear outlier in the A30P group, a Grubbs outlier test was conducted to statistically determine deviation from the group mean, accordingly 1 A30P mouse was excluded from analysis (see supplementary figure 1 for inclusive data). Correlative analysis was also performed between mean fluorescence for pS129 and γH2.AX within human cases via Spearman’s correlation (Prism). Additionally, frequency plots of immunoreactivity per channel were calculated, based on all sampled ROIs per disease condition or genotype as per NeuN +ve and NeuN - ve nuclei (Prism). Such quantification was done in order to visualise alterations in nuclear immunoreactivity at a cellular population level per disease condition or genotype.

### Quantification of genomic DNA damage derived chromatin in Lewy bodies

Sections from DLB cases, either stained with γH2A.X and pS129 aSyn or with 53BP1 (Rabbit anti-53BP1, 1:250, Cat# 4937S, Cell Signalling) and pS129 (staining protocol as outlined above) were imaged on a widefield microscope with a 20x objective lens. For the quantification of γH2A.X positive LBs, one section (from block I as outlined previously[54], mounted on a 76×39 mm slide, including BA21, 22, 41 and 42) from 10 DLB cases were visually inspected for all pS129 immunoreactive LB and co-localisation of γH2A.X determined. For all cases, no less than 25 LBs per section were sampled. Percentage LBs γH2A.X+ve and γH2A.X -ve were plotted as cumulative bar charts to establish a measure of the frequency of genomic DNA damage containing LBs within DLB cases. γH2A.X colocalization within LBs was further validated by confocal imaging at 63x oil immersion objective lens and Z-stack imaging.

### Subcellular fractionation of frozen post-mortem human tissue

Nuclear isolation was performed as previously reported [34]. In Brief, ∼250mg of frozen lateral temporal cortex from Con and DLB cases (n=9) were homogenised 1:16 (W:V) in nuclear extraction buffer (0.32 mM sucrose, 5 mM CaCl_2_, 3 mM Mg(Ac)_2_, 10 mM Tris-HCl, 0.1% NP-40, pH 8) supplemented with cOmplete proteases inhibitor cocktail and phostop tablets (1 per 10 ml, Sigma) via a dounce homogeniser (50 manual strokes per sample). Nuclei were then isolated from the resulting crude whole tissue homogenate via sucrose gradient centrifugation (1.8 M Sucrose, 3 mM Mg(Ac)_2_, 10 mM Tris-HCl, pH 8, 107,000 rcf x 2.5 hrs, 4°C). Nuclear pellets were washed twice in 0.01M PBS prior to being resuspended in 500 µl 0.01 M PBS, recovered and stored at −80°C prior to use.

### Western blot analysis

Nuclear fractions from Con (n=13) and DLB (n=12) were pretreated with DNAse (50 u/ml, 15 mins RT, RQ1 DNAse, Promega), were adjusted to 80 ng /µl with LDS sample buffer (Fisher Scientific), reducing agent (Fisher Scientific) and dH_2_O as per BCA protein concentration assay (Fisher Scientific). Prepared samples were heated at 70 °C for 10 mins and loaded at 4 µg/lane and separated via SDS page (4-12% Bis-Tris gels, Fischer Scientific) with MES buffer (200 V, 35 mins) and transferred to nitrocellulose membranes (0.2 µm) via Iblot 2 (7 mins, 20 mV, Fisher Scientific). Membranes were then washed in TBST, blocked in 5% milk powder (MP) containing TBST for 1 hr at RT on a rolling mixer, before being incubated in primary antibody (rabbit anti-γH2A.X, Cat#9718, Cell Signalling, 1:1000) containing 5% bovine serum albumin (BSA, Sigma) TBST, supplemented with 0.05% sodium azide, overnight at 4°C. Secondary antibody conjugation was performed with goat anti rabbit-HRP (1:5000, Sigma) in 5% MP containing TBST for 1hr at RT. Blots were washed in 3x TBST for 5mins in-between each incubation step. Immunoreactivity was visualised with enhanced chemiluminescence (1.25mM Luminol, 30 µM Coumaric acid and 0.015 % H_2_O_2_) and captured with a Fuji LAS 4000 and imaging software (Fuji LAS image, Raytek). Following image capture, membranes were re-stained for Histone H3 loading control (Mouse anti Histone H3, Cat# 3638S, Cell Signalling, 1:5000) and secondary antibody goat anti-mouse-HRP (1:5000, Sigma) as described above.

Immunoreactivity was measured using imageJ with area under curve function. γH2A.X signals were normalised to Histone H3 loading controls. Quantification of control and DLB nuclear isolates were performed across multiple western blots and as such values were normalised within blots to control values prior to being pooled and probed for significance via a Mann-Whitney non-parametric t-test.

## Mass spectrometry analyses of fractionated tissue

### Protein Digestion and quantitative proteomics

Frozen samples (250 mg) were fractionated as above, nuclear pellets were resuspended in 50µl of 5% SDS containing PBS. Proteins were digested using S-Trap (Protifi) and samples cleared of DNA/RNA by heating to 95 °C and sonication. Samples were reduced with TCEP (5mM final concentration, 15 min incubation at 55 °C) and alkylated with Iodoacetamide (10mM final concentration, 10 min incubation at RT). Resulting samples were acidified with 12% Phosphoric acid (final concentration of 2.5%, v/v), followed by addition of 6 vol. of loading buffer (90% methanol, 100 mM TEAB, pH 8) and loaded onto S-Trap cartridges. Cartridges were spun at 4000 x g for 30s and washed with 90% loading buffer x3 and flow-through discarded. Retained proteins were trypsin digested (10:1 protein: trypsin), in 50 mM TEAB, pH8.5 for 3h at 47°C.

Peptides were eluted, initially, with 50 µl of 5 0mM TEAB, followed by 50 µl of 0.2% formic acid and finally with 50 µl of 50% acetonitrile and 0.2% formic acid. Samples were then frozen and dried with centrifugal evaporator until at a volume of ∼1 µl. The sample was dissolved in 0.1 % Formic acid at a concentration of 1 µg/µl.

Peptide samples were loaded with 1 µg per LCMS run, peptides separated with a 125 min nonlinear gradient (3-40% B, 0.1% formic acid (Line A) and 80% acetonitrile, 0.1% formic acid (Line B)) using an UltiMate 3000 RSLCnano HPLC. Samples were loaded onto a 300μm x 5mm C18 PepMap C18 trap cartridge in 0.1% formic acid at 10 µl/min for 3 min and further separated on a 75 μm x 50 cm C18 column (Thermo EasySpray -C18 2 µm) with integrated emitter at 250 nl/min. The eluent was directed into a Thermo Orbitrap Exploris 480 mass spectrometer through the EasySpray source at a temperature of 320 °C, spray voltage 1900 V. The total LCMS run time was 150 min. Orbitrap full scan resolution was 60,000, RF lens 50%, Normalised ACG Target 300%. Precursors for MSMS were selected via a top 20 method. MIPS set to peptide, Intensity threshold 5.0 e3, charge state 2-7 and dynamic exclusion after 1 times for 35 s 10 ppm mass tolerance. ddMS2 scans were performed at 15000 resolution, HCD collision energy 27%, first mass 110 m/z, ACG Target Standard.

### Bioinformatic proteomic analysis

The acquired data was searched against the human protein sequence database, available from (https://www.uniprot.org/uniprot/?query=proteome:UP000005640), concatenated to the Common Repository for Adventitious Proteins v.2012.01.01 (cRAP, ftp://ftp.thegpm.org/fasta/cRAP), using MaxQuant v1.6.43. Fractions were assigned as appropriate. Parameters used: cysteine alkylation: iodoacetamide, digestion enzyme: trypsin, Parent Mass Error of 5ppm, fragment mass error of 10 ppm. The confidence cut-off representative to FDR < 0.01 was applied to the search result file. Data processing was performed in Perseus [74]. All common contaminants, reversed database hits and proteins quantified with less than 2 unique peptides were removed from the dataset. For quantitative proteomics, per case, intensity values for proteins were transformed to log2 and technical replicates averaged. Median was then subtracted within each sample to account for unequal loading and the width of the distribution adjusted. The dataset was filtered, keeping proteins with >2 valid values in at least one experimental group. Furthermore, any protein which was not detected in 70% of all samples were removed. Remaining missing values were imputed from the left tail of the normal distribution (2StDev away from the mean, +/-0.3 StDev). Compositional values were used to determine any changes in the abundance of identified proteins between Con and DLB nuclei samples.

Given that proteomic analysis was conducted in 3 batches (containing 3 control and 3 DLB cases, each). Values were transformed into Z-scores per protein per batch to allow for disease comparison, negating the impact of different runs across samples.

### Determination of altered expression and functional enrichment

Comparisons of nuclear protein abundance between disease groups were performed using Students T-test (S_0_ = 0.1, with permutation-based FDR<0.1, with 250 randomizations). Heatmaps of Z-scores per protein per case were plotted in Perseus. All proteins established as upregulated as per FDR<0.1, were subject to STRING database[70] and DAVID Bioinformatics[26, 65] analysis. For establishing enrichment of functional terms, upregulated proteins (81 proteins) were compared against a reference background dataset of all detected proteins (1810) in DAVID. Go terms (GOTERM_BP_Direct, GOTERM_CC_Direct and GOTERM_MF_Direct) and Functional annotations (UP_KW_BIOLOGICAL_PROCESS, UP_KW_CELLULAR_COMPONENT and UP_KW_MOLECULAR_FUNCTION) were used for assessment of enrichment as per the default setting of DAVID bioinformatics. Output clusters and enrichment data were inputted into R, generating graphs reporting -Log10 FDR and enrichment per GO term and/or keyword. Further determination of specific DNA damage/repair processes involving the identified upregulated proteins was perform via STRING. Here the reference set used was the human genome and not the background dataset, as this analysis was intended purely to highlight specific functional pathways which may be affected from changes in the nuclear proteosome.

## Results

### Elevation of double strand breaks in DLB cortical tissue and A30P mouse model of synucleinopathy

DNA damage markers of SSBs and DSBs were measured as per XRCC1 and γH2A.X in NeuN-positive (+ve) neurons and NeuN-negative (-ve) non-neuronal cells within the lateral temporal cortex of control and DLB cases (Fig 1.a). As expected immunoreactive signals of XRCC1 and γH2A.X were concentrated within nuclei. Mean levels of SSBs in both NeuN +ve and NeuN -ve nuclei were comparable between control and DLB cases (p>0.05, Fig 1.b). However, mean γH2A.X levels were elevated in NeuN +ve and NeuN -ve nuclei in DLB cases compared to controls (p<0.05, n=12 cases for both, Fig 1.c). Analysis of the frequency distribution of XRCC1 (Fig 1.d) and γH2A.X (Fig 1.e) intensities in all nuclei measured, demonstrated a γH2A.X selective rightward extension of the intensity profile for both NeuN +ve and NeuN -ve nuclei in DLB cases compared to controls. Such a change in the distribution profile of γH2A.X in DLB is consistent with an increase in nuclear DSBs across the cellular population, as opposed to being largely driven by a selective subpopulation of cells, such as those bearing LBs. There was no significant difference in the mean number of NeuN +ve and NeuN-ve cells sampled per field between controls and DLB cases (Fig1.f; p>0.05).

**Figure 1.**
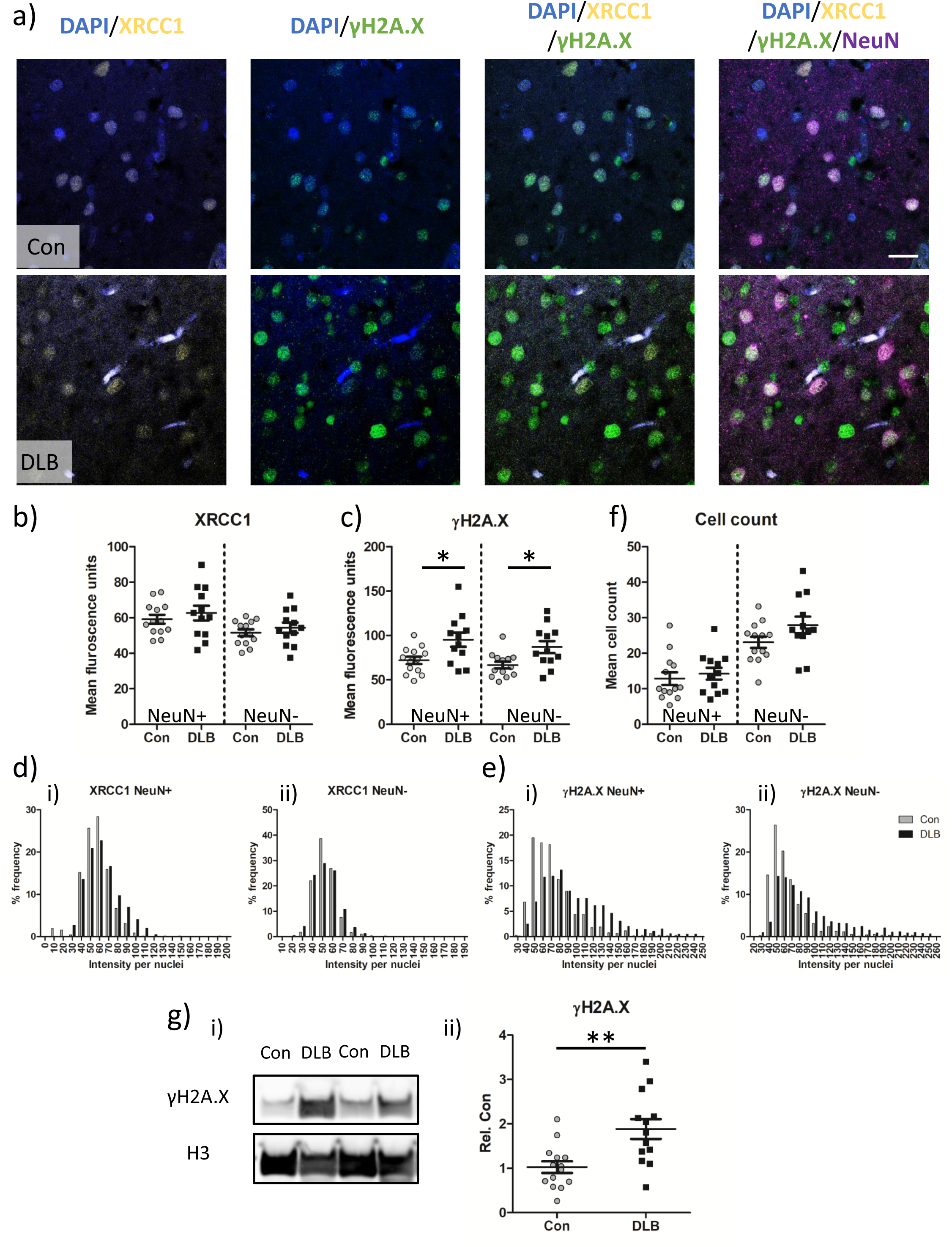
Double strand breaks are increased in temporal cortex nuclei of DLB cases. Example confocal images (x63 objective) from control (Con) and cases of dementia with Lewy bodies (DLB) showing single strand break (XRCC1) and double strand break (γH2A.X) associated immuno-fluorescence in neurons (NeuN) and non-neuronal cells. Nuclei are co-stained with DAPI (a). Quantification of mean XRCC1 (b) γH2A.X (C) fluorescence signal, alongside nuclear based cell counts (f), from Con (n=13 cases) and DLB (n=12 cases) are presented. Additionally, frequency distribution of XRCC1 (d.i-ii) and γH2A.X (e.i-ii) per nuclei are shown. Each measure output is reported for neuronal (NeuN+) and non-neuronal (NeuN-) populations. Western blot images of γH2A.X and loading control Histone H3 (g.i) and quantification of loading adjusted γH2A.X signal in Con and DLB cases (n=13 for both; g.ii) are also shown. Data are expressed in scatterplots with mean±SEM (in b,c,f and g) and as percentage frequency of immunofluorescence intensity per bins of 10 units width (in d + e). *=p<0.05 and **=p<0.01. Scale = 20 µm.

The elevation of nuclear γH2A.X signal in DLB cases compared to controls was confirmed via western blot of isolated nuclear lysates from frozen lateral temporal cortex (Fig 1.g, p<0.01, n=12 per group). Following the statistical identification of an outlier in the data as per Grubbs test (see supplementary figure 1), similar changes were observed in the somatosensory cortex of A30P mice at 2.5-3 months of age (Fig2a-c). However, in accordance with the neuronal specific expression of human mutant aSyn in these mice, levels of the DSB marker γH2A.X were only elevated in the NeuN +ve neuronal population (p<0.05, Fig 2.b) and not in non-neuronal nuclei (p>0.05). Notably, despite no robust change in the mean values of SSB marker XRCC1 between A30P and WT mice, there was an apparent shift in the frequency distribution of XRRC1, as well as in the distribution of γH2A.X in neuronal NeuN +ve and non-neuronal NeuN -ve nuclei (Fig 2 .d+e). The rightward shift indicative of at increased percentage of nuclei with high levels of both SSBs and DSBs within the somatosensory cortex, as a consequence of the expression of mutant aSyn. Despite the elevation of DNA damage in these young A30P mice there was no evidence of cell loss, as neuronal and non-neuronal numbers were comparable between WT and A30P mice (fig 2.f, p>0.05).

**Figure 2.**
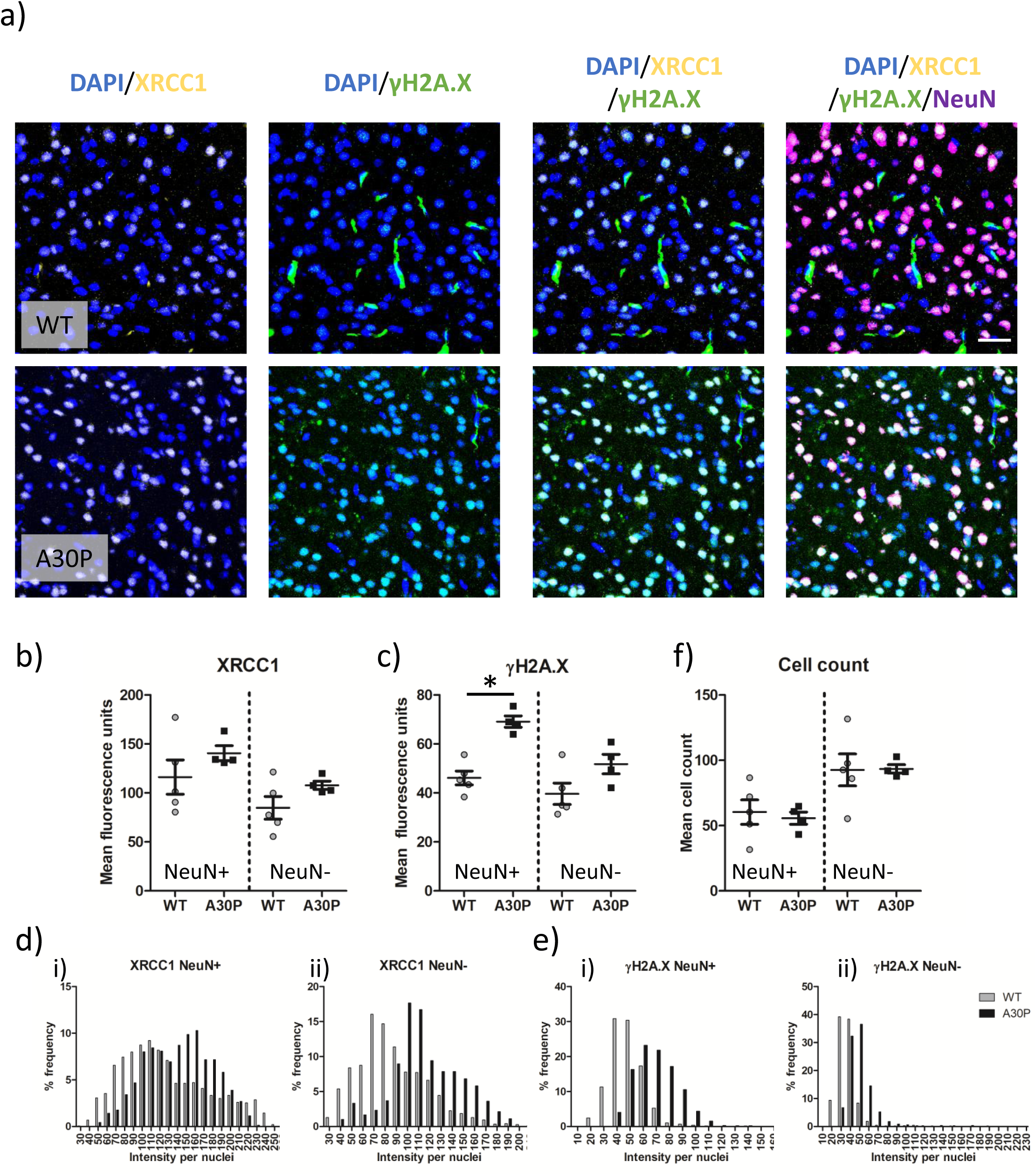
Assessment of single and double strand breaks in an A30P mouse model of synucleinopathy. Immunofluorescence confocal images (63x objective lens) of XRCC1, γH2A.X, NeuN and nuclei co-stained with DAPI from the somatosensory cortex of wild type (WT) and A30P mice (a). Quantification of neuronal (NeuN +) and non-neuronal (NeuN -) mean XRCC1 (b) γH2A.X (c) signal in WT (n=5) and A30P mice (n=4). Frequency distribution of XRCC1(d) and γH2A.X (e) per nuclei are shown in NeuN + (i) and NeuN – (ii) nuclei are also shown alongside cell count, as per nuclei number (f). Data are expressed in scatterplots with mean±SEM (in b, c and f) and as percentage of immunofluorescence intensity binned with 10 unit widths (d and e), * =p<0.05. Scale= 20 µm.

### Pathological nuclear aSyn correlates with an increase in DSBs

To determine whether nuclear aSyn pathology correlated with the observed increase in DNA damage, levels of phosphorylated nuclear aSyn (pS129) and DSBs as per γH2A.X were quantified via immunostaining. Consistent with the initial observations (Fig 1.), DSBs were again observed as increased in both NeuN +ve and NeuN -ve nuclear populations (Fig 3. a+b). As previously reported[34], nuclear pS129 aSyn was increased in the DLB cohort, in NeuN +ve and NeuN -ve nuclei (p<0.01, for both, Fig 3. a+c). Similarly, a rightward shift in the distribution profile of mean intensity of nuclear pS129 aSyn was seen in NeuN +ve and NeuN-ve nuclei populations (Fig 3.d i + ii), indicative of a general rise in nuclear pS129aSyn at the cellular population level. Correlative analysis per case of the mean intensity of γH2A.X and nuclear pS129 aSyn, reported significant linear relationships in both NeuN +ve (Fig 3.e.i, r=0.49, p<0.05) and NeuN -ve (Fig 3.e.ii, r=0.55, p<0.01) nuclei, supportive of a potential interaction between nuclear aSyn pathology and elevated DSBs.

**Figure 3.**
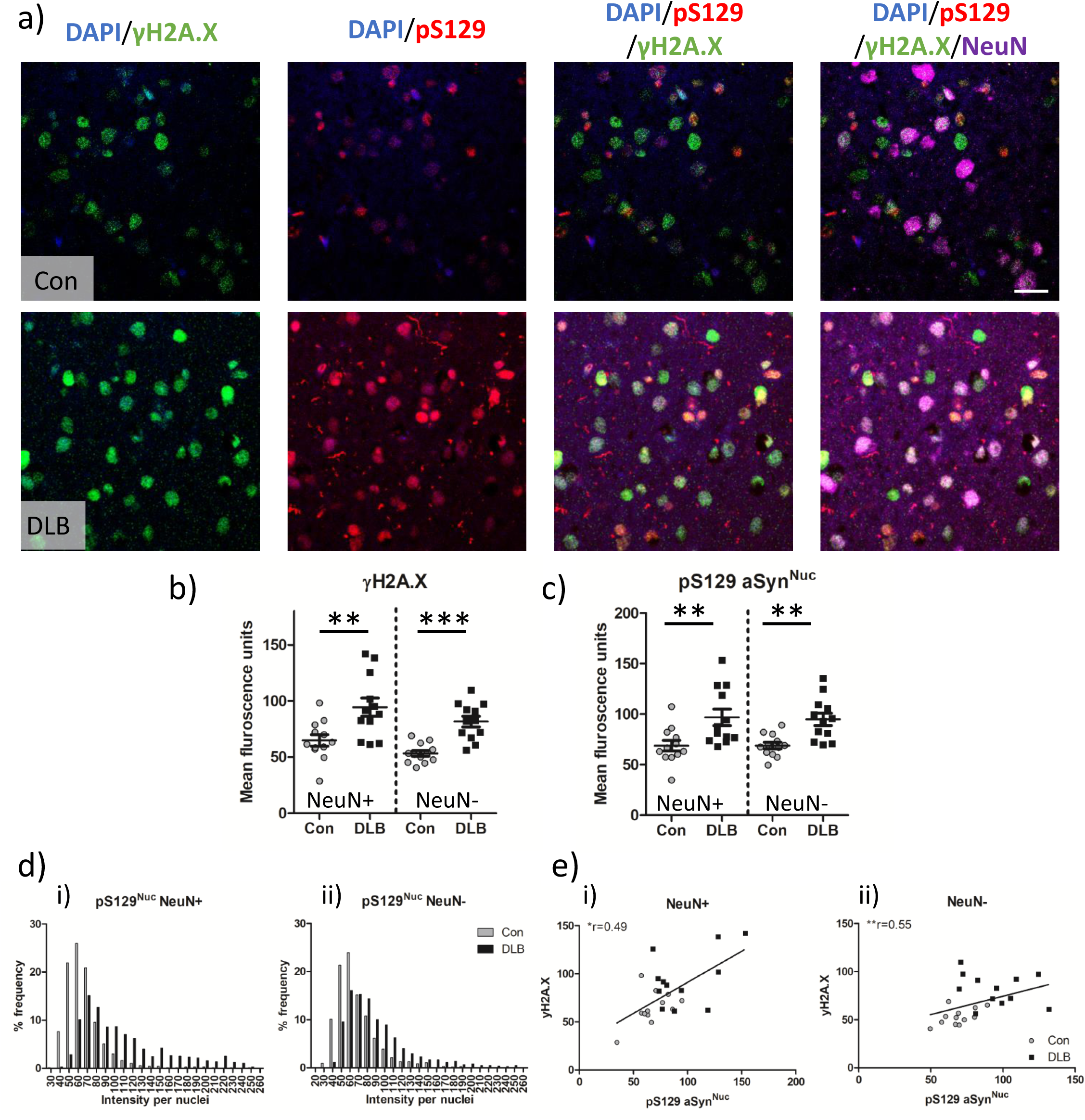
Phosphorylated aSyn increases in concert with double strand breaks and correlate with each other in the nuclear compartment. Example confocal images (x63 objective) from control (Con) and cases of dementia with Lewy bodies (DLB) showing alpha-synuclein phosphorylated at serine 129 (pS129), double strand break (γH2A.X) immuno-fluorescence in neurons (NeuN) and non-neuronal cells. Nuclei are co-stained with DAPI (a). Quantification of nuclear pS129 (pS129 aSyn^Nuc^) (b) and γH2A.X (c) fluorescence from Con (n=13 cases) and DLB (n=12 cases) are presented. Frequency distribution of ps129 (d.i-ii) per nuclei are shown as correlative levels of ps129 aSyn^Nuc^ with levels of γH2A.X, per case, with spearman’s correlation (r) reported (e.i-ii). Measures are reported for neuronal (NeuN+) and non-neuronal (NeuN-) populations. Data are expressed in scatterplots with mean±SEM (in b,c,e) and as percentage frequency of immunofluorescence intensity per bins of 10 units width (in d). *=p<0.05, **=p<0.01 and **=p<0.001.

### LBs contain genomic chromatin originated from DSBs

Remarkably, the DSB marker γH2A.X was not only present in the nuclei of cortical cells in fixed tissue sections but was also apparent in pS129-aSyn rich LBs within DLB cases (Fig 4.a). In comparison to the robustly labelled γH2A.X LBs (white arrow heads), Lewy neurites (white arrows) were negative for the DSB marker. Despite the prominence of this nuclear DSB marker, LBs were negative for a secondary nuclear marker 53BP-1 (Fig 4.b). Such an immunoreactivity profile is consistent with the genomic chromatin originating from cytoplasmic chromatin fragments, downstream from excessive unrepaired DNA damage[44]. γH2A.X immunoreactivity within LBs was confirmed via confocal microscope, such that γH2A.X puncta were evident throughout the Lewy body structure (Fig 4.c). Intriguingly, in addition to a high level of γH2A.X within LBs, several examples of perinuclear aSyn accumulations, assumed to be precursors of LBs[48], were also positive for γH2A.X (Fig 4.d). Across 10 DLB cases, a remarkably high percentage of LBs, as assessed from one section per case, were positive for γH2A.X (Fig 4.e, mean 89.6 ± 4.2 %). Such frequency of γH2A.X immunoreactivity was largely consistent across cases (9 of 10 cases, with 90-100% of LBs positive for γH2A.X, with one case of lower frequency of 52.9%), suggesting that the inclusion of DSB originating genomic DNA within LBs is a common occurrence.

**Figure 4.**
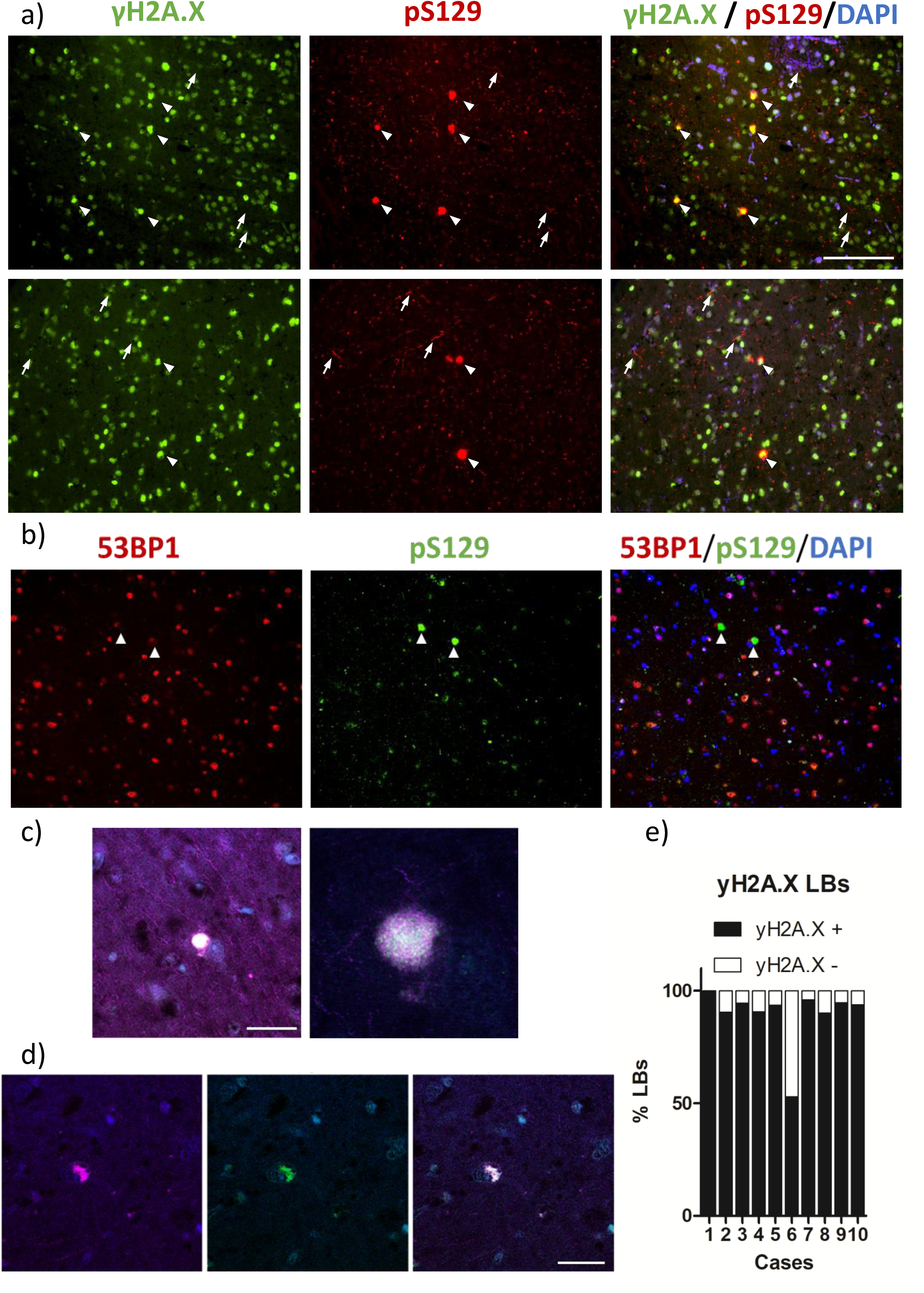
Nuclear material is a core constituent of cortical Lewy bodies. Example widefield images (x20 objective) cases of dementia with Lewy bodies (DLB) showing Lewy bodies (LBs) as detected via alpha-synuclein phosphorylated at serine 129 (pS129), *γ*H2.AX (a) and 53BP-1 (b) immuno-fluorescence, with nuclei are co-stained with DAPI. Note the colocalization of γH2.AX with extranuclear pS129 Lewy bodies (a; white arrow heads) and absence of colocalization of 53BP-1 with Lewy bodies (b; white arrow heads). Confocal image of perinuclear rounded LB (c) and peri nuclear aSyn accumulation (d) with pS129-aSyn and γH2A.X colocalised. Quantification of γH2.AX positive (γH2A.X +) and negative (γH2A.X -) LBs in ten examined cases (e). Data shown as percentage LBs positive and negative for γH2A.X.

### Upregulated DNA damage repair proteins in the nuclear proteome of DLB cortical tissue

Given the widespread elevation of DSBs within DLB cortical tissue and the potential involvement of DSB related genomic material in the cytoplasmic aggregation of aSyn, a key aspect to the cellular pathology underlying DLB is likely to be disease dependent changes in the nuclear proteome.

Proteomic analysis of nuclei, isolated from the lateral temporal cortex of DLB and control tissue (n=9 for each), reported 1810 identified proteins detected consistently across disease conditions, 81 (4.5%) of which were upregulated and 6 (0.3%) of which were downregulated in DLB tissue (Fig 5.a +b, for full data set see supplemental table 2). No protein was reliably detected only in either control or DLB cases.

**Figure 5.**
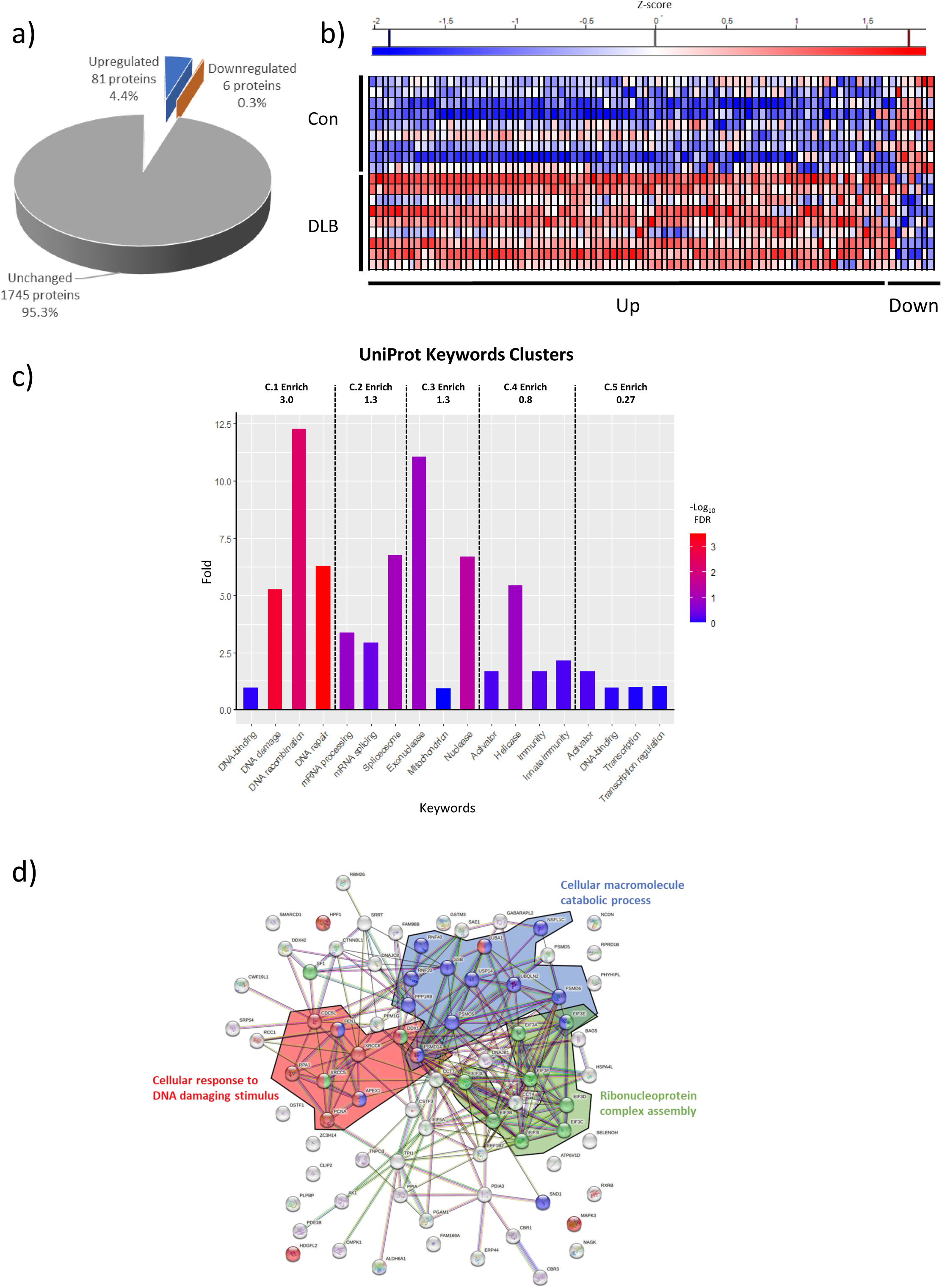
Altered nuclear proteosome of DLB temporal cortex tissue. Upregulated, downregulated and unchanged nuclear proteins in temporal cortex of dementia with Lewy body cases (DLB), compared to controls (Con). Data provided as absolute numbers and percentages of total identified proteins (a). Z-score heatmap of altered proteins between Con and DLBs (b). Uniprot keyword enrichment of the 83 proteins established as elevated in nuclear proteosome of DLB compared to controls. Fold enrichment and significance is shown as per false discovery rate for each individual keyword. Keywords are grouped into clusters as reported per the DAVID bioinformatic database. Dotted lines denote the boundaries of clusters, with cluster enrichment (C.Enrich) from background reference database shown within each cluster (c). String map of protein interactome of upregulated proteins. Three prominent functional categories are highlighted, Cellular response to DNA damaging stimulus (red), Cellular macromolecule catabolic process (blue) and Ribonucleoprotein complex assembly (green), with individual proteins associated with each term denoted by colour and hubs of strong interaction associated with each term highlighted by background colour (d).

Functional cluster analysis against Uniprot keywords as per DAVID bioinformatics reported a primary enrichment cluster (3 fold enriched) of DNA binding, damage, recombination and repair, with a secondary cluster (1.3 fold enriched) associated with mRNA processing and splicing as well as a third cluster (1.3 fold enriched) associated with nuclease activity (Fig 5.c + supplementary table 3). Similarly, analysis of upregulated proteins for enriched GO terms reported a cluster (1.2 fold enriched) associated with DNA processing, base-excision repair and binding of damaged DNA although individual terms were not reported as significantly enriched following adjustments for multiple comparisons (See supplementary table 4). Regardless, from this GO term analysis, clusters associated with Ribonucleoprotein complex assembly (4.1 fold enhanced) and Proteasome catabolic processing (1.2 fold enhanced) were also reported (see Supplemental figure 2 + supplementary table 4). Given that only 6 proteins were found to be reduced in the DLB nuclei compared to controls, no functional enrichment was conducted (See Supplementary table 2 for protein list).

To further understand the interactions of the upregulated proteins, additional STRING database analysis was conducted (Figure 5.d). From the 81 proteins, 79 were mapped within the database and reported a strong level of interaction (79 nodes, 238 edges, average node degree 6.03, PPI enrichment p<1.0e^-16^). Visual inspection of the String network demonstrated three prominent biological process clusters: Cellular response to DNA damaging stimulus (GO:0006974), Cellular macromolecule catabolic processes (GO:0006074) and Ribonucleoprotein complex assembly (GO:0022618).

### Upregulated base excision and double strand break repair pathways in DLB cortical tissue

Those proteins associated with “Cellular response to DNA damaging stimulus” demonstrated a largely consistent pattern of upregulation across DLB cases compared to controls (Fig 6.a). When resubmitted to the STRING database for protein interaction analysis (Fig 6.b), unsurprisingly these proteins demonstrated as strong protein-protein interaction (13 nodes, 21 edges, average node degree 3.23, PPI enrichment p=1.53e^-06^) and key biological functions including “Base-excision Repair” and “Double strand break repair via non-homologous end-joining” were reported (See table 2 for top 5 terms, based on strength, and Supplementary table 5 for full list and supplementary table 6 for functional assignment).

**Figure 6.**
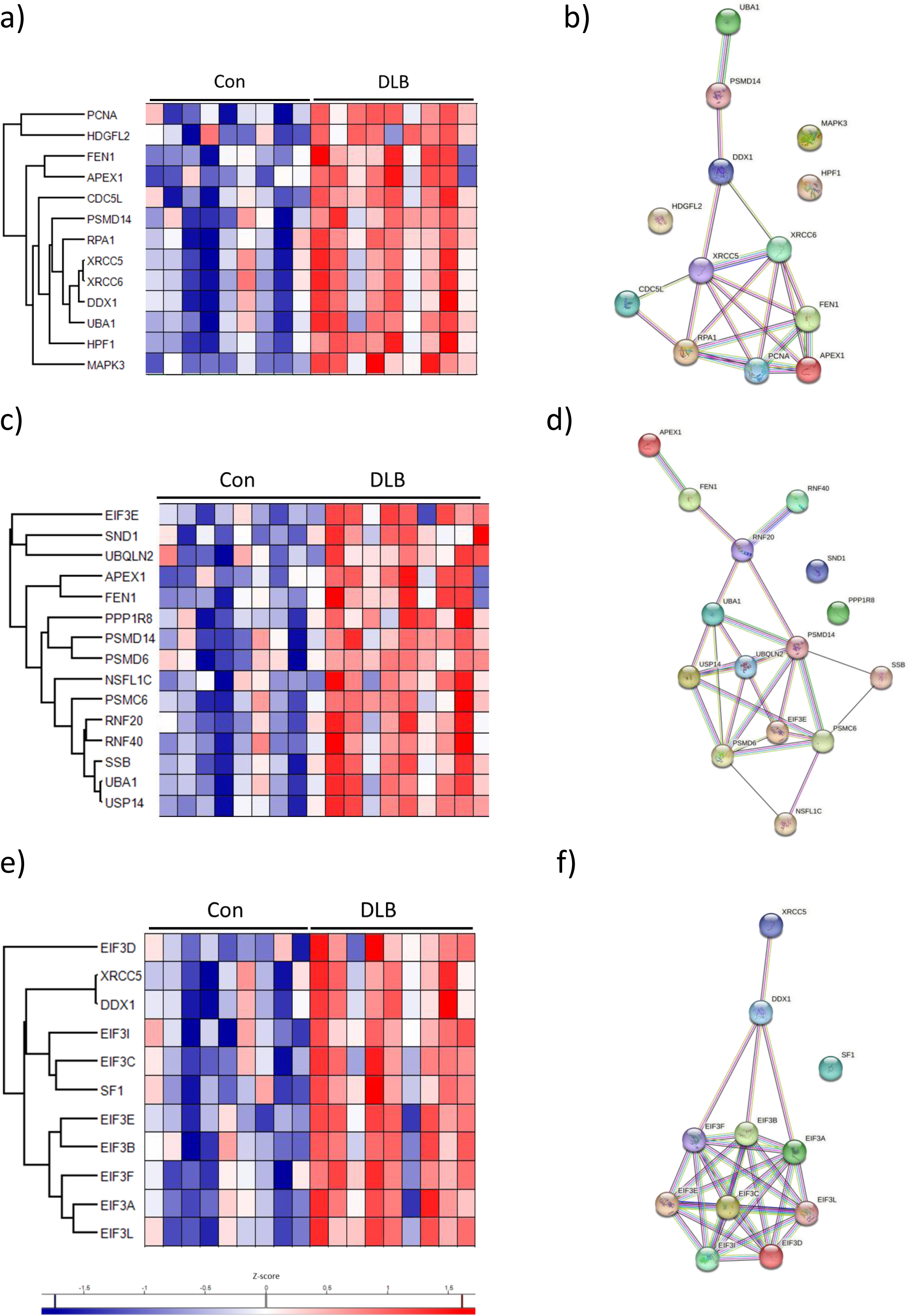
Interaction mapping of upregulated string functional analysis in the nuclear proteosome of DLB cortical tissue. Z-score heatmap of identified proteins associated with the cellular response to DNA damage proteins (a), cellular macromolecule catabolic process (c) and ribonucleoprotein complex assembly (e) organised as per individual cases of either controls (Cons) or cases of Dementia with Lewy bodies (DLBs). Proteins are arranged per Euclidean distance between Z-scores. String map of proteins are organised as presented per functional category (b,d and f).

**Table 2.** STRING biological processes GO terms associated with enriched protein groups. Top 5 GO terms, based on enrichment strength. List is subdivided into cellular response to DNA damage, Cellular metabolic catabolic process, and Ribonucleoprotein complex assembly. Provided are the GO term identification number (Go Term ID), Term description, observed protein number within the list, background protein number, strength of enrichment, FDR adjusted p-value and proteins identified.

| GO Term ID | Term description | observed | background | strength | FDR | Proteins involved |
| --- | --- | --- | --- | --- | --- | --- |
| **Cellular response to DNA damage** |  |  |  |  |  |  |
| GO:0032201 | Telomere maintenance via semi-conservative replication | 2 | 7 | 2.64 | 0.0069 | FEN1,PCNA |
| GO:0006287 | Base-excision repair, gap-filling | 3 | 14 | 2.51 | 0.00012 | APEX1,FEN1,PCNA |
| GO:0071481 | Cellular response to X-ray | 2 | 12 | 2.4 | 0.0158 | XRCC6,XRCC5 |
| GO:0006284 | Base-excision repair | 4 | 43 | 2.15 | 1.74e-05 | APEX1,RPA1,FEN1,PCNA |
| GO:0006303 | Double-strand break repair via nonhomologous end joining | 3 | 43 | 2.02 | 0.0018 | XRCC6,XRCC5,PSMD14 |
| **Cellular metabolic catabolic process** |  |  |  |  |  |  |
| GO:0071894 | Histone H2B conserved C-terminal lysine ubiquitination | 2 | 3 | 2.94 | 0.0027 | RNF40,RNF20 |
| GO:1903069 | Regulation of ER-associated ubiquitin-dependent protein catabolic process | 2 | 10 | 2.42 | 0.0164 | USP14,UBQLN2 |
| GO:0006287 | Base-excision repair, gap-filling | 2 | 14 | 2.27 | 0.0266 | APEX1,FEN1 |
| GO:0006401 | RNA catabolic process | 5 | 164 | 1.6 | 9.59e-05 | EIF3E,FEN1,PPP1R8,SND1,SSB |
| GO:0061136 | Regulation of proteasomal protein catabolic process | 5 | 196 | 1.53 | 0.00017 | USP14,RNF40,UBQLN2,PSMD14,PSMC6 |
| **Ribonucleoprotein complex assembly** |  |  |  |  |  |  |
| GO:0075525 | Viral translational termination-reinitiation | 4 | 5 | 3.16 | 6.47e-09 | EIF3D,EIF3B,EIF3A,EIF3L |
| GO:0001732 | Formation of cytoplasmic translation initiation complex | 8 | 16 | 2.95 | 2.99e-18 | EIF3D,EIF3E,EIF3C,EIF3B,EIF3A,EIF3I,EIF3F,EIF3L |
| GO:1902416 | Positive regulation of mRNA binding | 3 | 8 | 2.83 | 1.11e-05 | EIF3D,EIF3E,EIF3C |
| GO:0075522 | IRES-dependent viral translational initiation | 4 | 11 | 2.81 | 5.60e-08 | EIF3D,EIF3B,EIF3A,EIF3F |
| GO:0019081 | Viral translation | 5 | 15 | 2.78 | 4.52e-10 | EIF3D,EIF3B,EIF3A,EIF3F,EIF3L |

A consistent pattern of upregulated expression and strong protein-protein interactions (Fig 6. C + d ; 15 nodes, 21 edges, average node degree 3.47, PPI enrichment p= 2.61e^-13^) was also observed for those proteins associated with “Cellular macromolecule catabolic processing”. Key biological functions included “regulation of proteasomal protein catabolic process” but also “Base-excision repair-gap filling”, “RNA catabolic processing” and “Histone H2B conserved C-terminal lysine ubiquitination”, functions which are also associated with DNA damage and repair (see table 2 and Supplementary table 6 + 7).

Similarly, those proteins associated with “Ribonucleoprotein complex assembly” were consistent in their upregulation in DLB cases compared to controls (Fig 6.e) and also had robust protein-protein interactions (Fig 6.f; 11 Nodes, 32 edges, average node degree 5.82, PPI enrichment p< 1.0e^-16^). Key biological functions of these proteins were largely reported as related to protein translation, without overt association to DNA damage and repair (although, see table 2 and Supplementary table 6 + 8), see discussion for further details.

## Discussion

In the present study, widespread DSB accumulation was evident in the lateral temporal cortex of DLB cases and similar observations made early in pre-symptomatic A30P aSyn expressing mice. DSBs coincided with increased phosphorylated nuclear aSyn in DLB cases alongside the striking occurrence of γH2A.X within LBs and perinuclear aSyn aggregates, suggestive of a reciprocal relationship between aSyn pathology and genomic damage. Accordingly, we observed a robust upregulation of DNA damage and repair proteins, as well as ubiquitin mediated catabolic processing and the ribonucleoprotein complex within the nuclear proteosome of DLB temporal cortex. Collectively the data suggests an association of nuclear aSyn pathology with genomic DNA damage which in turn is associated with cytoplasmic aSyn pathology.

### Nuclear aSyn pathology and DNA damage

Building on our prior studies of the nuclear localisation of aSyn [23, 34, 53] and its pathological modification in DLB cortical tissue[34], we here demonstrate a disease dependent accumulation of DSBs as per γH2A.X in neuronal and non-neuronal cells in the temporal cortex of DLB cases. γH2A.X results from the rapid serine 139 phosphorylation of histone H2.A and is instrumental in the initiation of DSB repair, facilitating DDR molecule recruitment and is terminated upon repair completion [39]. Accordingly, increased γH2A.X suggests an increase in DSB occurrence and/or a diminished capacity for DSB repair.

Similar reports of increased DSBs in PD cases[6, 24], demonstrate a robust link between synuclein pathology and DNA damage, irrespective of Lewy body disease (LBD) subtype. However, unlike observed elevations of SSBs in PD brains [24], we did not detect an increase in SSBs in our DLB cohort. In post-mitotic cells, DSBs result from various cellular events, including close-proximity SSBs and exposure to external toxins or radiation[7]. Assuming the absence of exogenous agents, SSBs remain the most likely source of DSBs, yet as no overt increase in SSBs were evident, our findings are compatible with the notion that deficits within DSB repair mechanisms likely drive DSBs accumulation (although see discussion later). Accordingly decreased DSB repair rates were noted in aSyn knockout mice [57] and fibroblasts from A53T mice[79]. Moreover, the accumulation of DSBs have been reported in several model systems utilizing aSyn overexpression [53, 80] and pre-formed fibrils [25]. These studies together suggest that pathological aSyn accumulation or modification may mimic a loss of physiological function in DNA repair.

Indeed, it has been suggested that aSyn aggregation may sequester nuclear aSyn from its repair function, and as such a modest association between aSyn aggregates and higher levels of DSBs within specific neurons have been reported [57]. However, our analysis of DSBs here, reveals widespread elevation across neuronal and non-neuronal cells, extending beyond LB burdened neurons. Likewise, elevated DSBs in A30P mice at a young age (3m), preceding robust cytoplasmic aSyn accumulation and symptomatic presentation [58], implicates DNA damage repair dysfunction as an early occurring cellular insult, independent from mature aSyn aggregation.

In the present study, the extent of DSBs within a given case correlated with nuclear pS129-aSyn levels. pS129-aSyn is a prominent pathological hallmark of synucleinopathies and is widespread throughout PD and DLB brain tissue [55]. Similarly, elevations of pS129-aSyn occur early within genetic aSyn mouse models, including in A30P mice, where robust nuclear pS129 aSyn is amongst the earliest pathological changes [59]. The coincidence of nuclear pS129 and DSBs in both human disease tissue and in mouse models is indicative of the potential for aSyn phosphorylation to alter its modulation of DDR. This is consistent with the differential DNA-binding behaviour of pS129 aSyn compared to non-phosphorylated aSyn [15, 57] and the phospho-regulation of aSyn nuclear import [18, 23, 53]. In itself, increased nuclear aSyn can perturb nuclear functions as directed nuclear aSyn localisation also results in DNA damage accumulation [21, 51–53]. Moreover, in the absence of aSyn aggregation, the overexpression of nuclear aSyn can recapitulate PD based neurodegeneration and symptomology in mice [21]. Together this raises the possibility that independent of LB formation, cellular dysfunctions may be driven by the loss of a physiological aSyn nuclear function associated with DNA damage and its repair.

### Presence of DSB in LBs

Strikingly, this study found that the principal marker of DSBs, γH2A.X was also evident within the majority of cortical LBs examined as well as within perinuclear aSyn aggregates, hypothesized to be precursors of mature LBs[48]. Several studies have highlighted the presence of DNA within the core of LBs, although this has largely been assumed to be mitochondrial in origin Moors, 2021 #20;Power, 2017 #19}. Here, the histone marker yH2A.X, demonstrates that such DNA is also of nuclear origin. The occurrence of ectopic cytoplasmic genomic material (eCGM), either in the form of cytoplasmic chromatin fragments (CCFs) or micronuclei is a known consequence of chronic unresolved DNA damage [44]. Given that double stranded DNA Cherny, 2004 #27} and histones [22], the major components of chromatin, have pro-aggregation properties towards aSyn, our observations here, suggests that genomic DNA damage mediated eCGM may act as co-aggregates facilitating the formation of aSyn aggregates in-situ.

### Alteration of Nuclear DNA damage repair pathways

Concordant with widespread DSB accumulation, nuclear proteomic analysis highlighted an upregulation of DNA damage, and repair associated proteins. This is in contrast, to our prior observations of downregulated DDR related transcripts following aSyn overexpression in cell lines [51, 53]. Whilst mechanistic differences between aSyn overexpression and idiopathic synucleiopathy induced DNA damage may be a confounding factor, it must also be considered that transcript changes may not reflect alterations at a protein level[78]. Equally, our current post-mortem observations may be influenced by survivor bias. Nevertheless, such a premise would still imply a major role for DNA damage in LBD pathophysiology. Regardless, two major DNA damage repair pathways were highlighted via enrichment analysis, BER and DSB-repair via NHEJ with additional DDR components associated with DSB repair via HR also identified (Fig. 7+ 8 and s. table 6).

**Figure 7.**
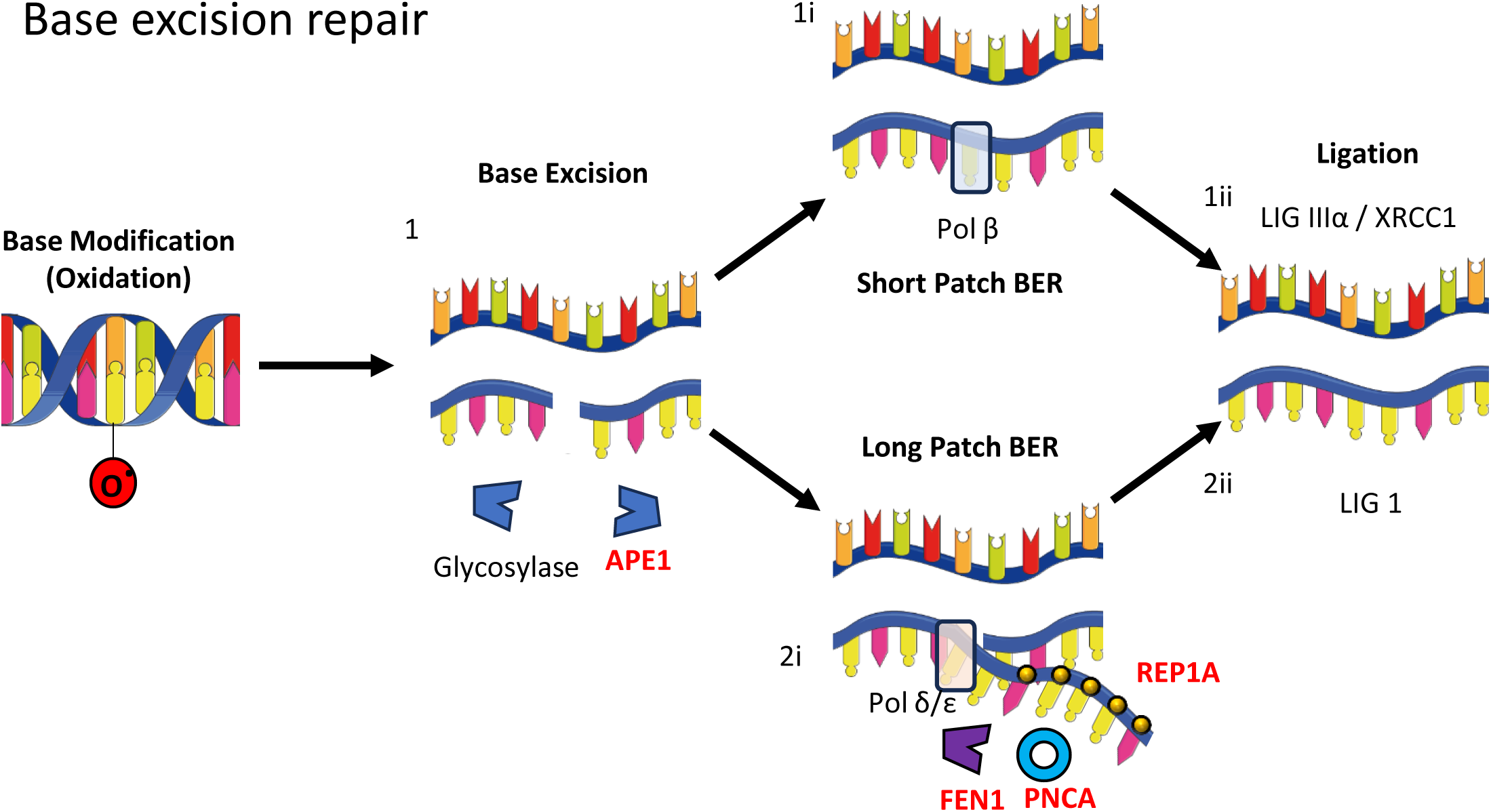
Base excision repair and upregulated protein components in DLB. Diagram of base excision repair (BER) pathways (a). Upon the oxidation (as shown) or another modification such as alkylation, deamination, modified bases are excised via the sequential processing of a damage specific glycosylase and APEX1 (1) and repair proceeds as per short patch (i) or long patch repair (ii). In short patch BER, the single nucleotide gap is repaired with the correct base by β-polymerase (2.i) and ligated by a DNA ligase IIIα-XRCC1 complex (2.ii). In cases of excess oxidation, long patch BER, a 2-8 nucleotide sequence is generated by δ/ε polymerases within the excision gap, this invaded single strand DNA is bound to replication protein 1 A (RPA1A), prior to the cleavage of the excessive sequence via the FEN1 endonuclease, proliferating nuclear protein A (PCNA) facilitating the process (2.ii) before being ligated by DNA ligase I [14, 27] (3.ii). BER proteins established as upregulated in DLB nuclei are indicated in red.

### Base excision repair

BER is activated as a consequence of alkylative, deaminative and oxidative base modification [30]. Driven by damage specific glycosylases and the endonuclease activity of apurinic/apyrimidine endodeoxyribonuclease 1 (APEX1 /APE1), BER proceeds via short or long patch repair (Figure 7). Alterations to BER as a consequence of synucleiopathy have previously been reported, including an elevation of abasic sites [56] and key BER nucleases [2, 20, 45, 67]. Here, upregulation of principle BER nucleases APE1 and FLAP endonuclease 1 (FEN1) as well as other regulatory proteins, is consistent with the overactivation of long patch BER. In line with the prominent involvement of oxidative stress in LBDs [12, 16] including the accumulation of oxidised and abasic mitochondrial DNA bases [1, 56, 82], our work implies that similar oxidative damage may occur within the nucleus of DLB cases. Engagement of BER may indeed be a protective response, as observed in oxidative stress models [75]. However, prediction of any beneficial repair capacity enhancement following DDR protein upregulation is difficult, as such isolated overexpression can often be detrimental to DNA damaging stressors and lifespan [63]. Indeed, excessive long patch BER activation can, as a consequence of excess nuclease activity, result in DSB formation rather than base repair [17, 64, 66, 68, 77]. Thus, the potential exists for BER overactivation in response to DLB pathology to contribute to the accumulation of DSBs.

### Double strand break repair

NHEJ is the principal pathway of DSB repair for post-mitotic neurons. NHEJ is conducted independent of a DNA repair template, with successive ligation and nucleotide addition often resulting in erroneous additions and deletions in the genetic code [9]. This in itself may drive increased rates of somatic mutations reported in the central nervous system of LBD cases [33]. NHEJ is conducted as opposed to HR, an error-less, cell cycle dependent DNA template driven repair mechanism [60], typically not employed by neurons but may be engaged in glial cells. Whilst sharing several common modulators, NHEJ and HR largely diverge in the molecular components required for successful DDR, both NHEJ and HR mediators being upregulated in DLB cortical nuclei (Figure 8). Thus, the data suggests at least a partial overactivation of both NHEJ and HR in DLB.

**Figure 8.**
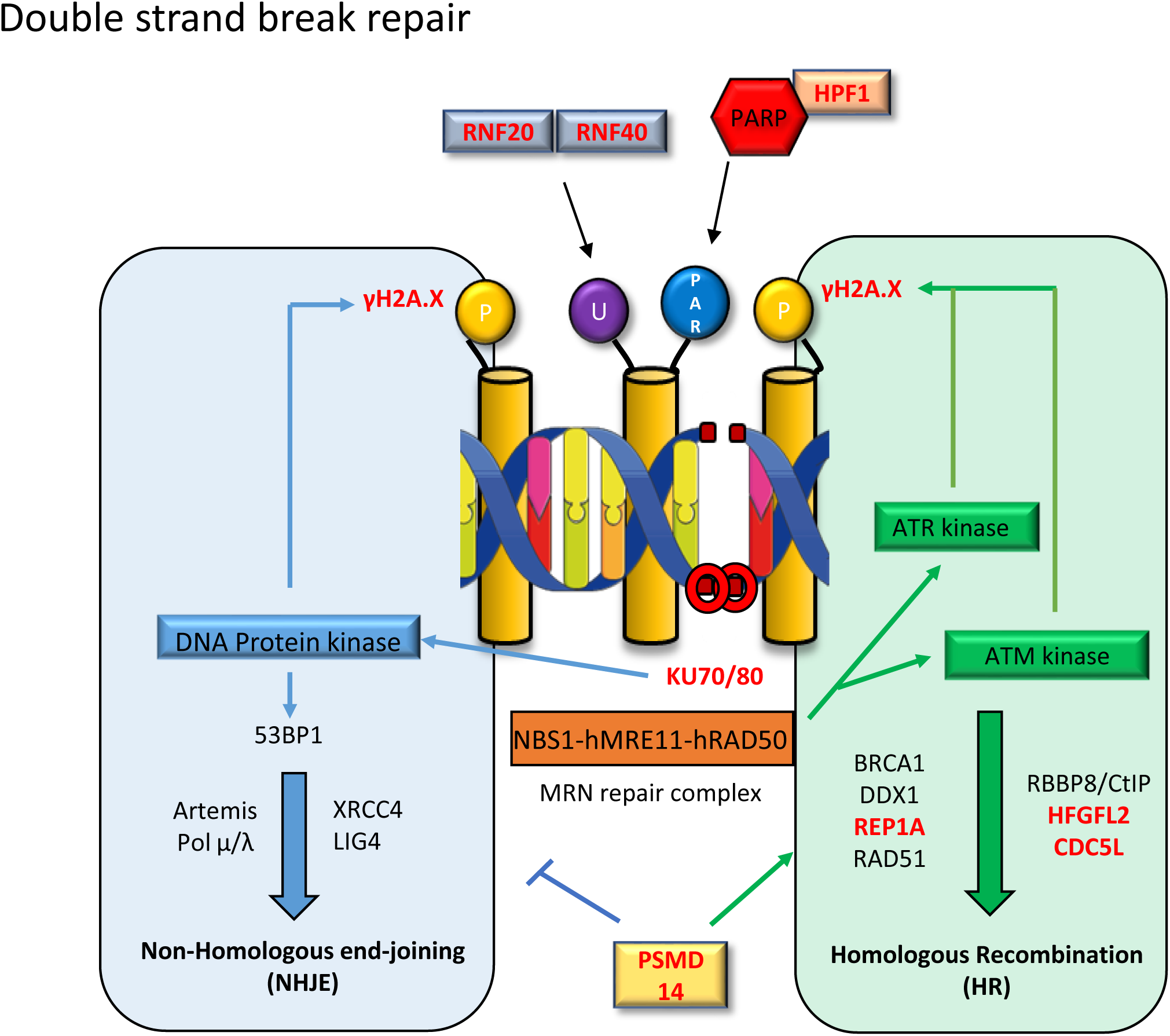
Double strand break repair pathways and upregulated protein components in DLB. Diagram of double strand break (DSBs) repair pathways, non-homologous end-joining (NHEJ) and homologous recombination (HR) are shown. In NHEJ, DSBs are sensed by DNA-protein kinase (DNA-PK) regulatory subunits KU70/KU80, in turn activating DNA-PK, enabling the phosphorylation of many substrates including histones, consequently generating DSB signal γH2A.X. To enable the recruitment of NHEJ machinery, chromatin surrounding the DSB is relaxed, in part due to histone ubiquitination (U) via ubiquitin ligases including ring finger 20/40 (RNF20/40)[69]. With accessibility to the DSB increased NHEJ components are recruited to the DSB site, including the NHEJ facilitatory 53BP1, Artemis nuclease for strand resection, DNA polymerase µ/λ for DNA synthesis and XRCC4/DNA ligase 4 for repair ligation[43]. In contrast, HR is initiated via the sensing of DSBs by MRN repair complex, activating Ataxia-telangiectasia mutated (ATM) and ataxia telangiectasia and Rad3-related (ATR) protein kinase, leading to the phosphorylation of many DDR substrates including the generation of γH2A.X[10]. CDC5L is a regulator of ATR kinases [83]. DSBs strands are resected by MRE11 promoted by BRCA1 as well as RBBP8/CtIP [10], which is facilitated by HDGFL2[3], creating 3’ overhangs, which are cleared of RNA-DNA hybrids via DEAD Box 1 (DDX1)[35, 36], prior to REP-A coverage and RAD51 guided homologous DNA search and strand invasions prior to the formation of holiday junction and repair[10]. Engagement of HR as opposed to NHEJ is in part co-ordinated by PSMD14, which restricts the accumulation of NHEJ inducer 53BP1 at damaged sites[31]. DSB repair protein upregulated in DLB nuclei are indicated in red.

Interestingly X-ray complementing repair cross components 5/6 (XRCC5/6 / Ku70 and Ku80), initial mediators of NHEJ and regulatory subunits for the DNA-protein kinase (DNA-PK) [43] were amongst the altered DDR proteins. Ku70/80 have previously been reported to be upregulated in a cell model of aSyn overexpression alongside the induction of DSBs [80]. Whilst intuitively, increased Ku70/80 could be anticipated as beneficial to the DSB repair capacity of cells, in rodent fibroblasts, DNA-PK activity was diminished and repair rates slowed, following the overexpression of the regulatory subunits [32]. Likewise in C9Orf72 expansion repeat ALS patient stem cell derived neurons, knockdown of upregulated Ku70/80 was reported as protective [38]. Thus, functional studies are now required to establish the impact of this upregulation in the context of aSyn pathology.

### Upregulation of ubiquitin regulators and eukaryotic initiation factor complex 3 proteins

Alongside the upregulation of DDR, cellular metabolic catabolic processes specifically involved in the ubiquitin/proteasome system (UPS) were also reported as upregulated in DLB cortical nuclei. The UPS is a critical regulator of DDR, regulating chromatin relaxation, protein-protein interactions essential for DDR as well as more conventionally mediated proteasomal degradation of DDR effectors and substrates [[8, 49, 69]. Accordingly, UPS upregulation is consistent with a nuclear environment in which DNA damage, repair and it’s associated histone turnover is highly engaged. Finally, proteins associated with the ribonucleoprotein complex assembly, specifically eukaryotic initiation factor complex 3 (EIF3) proteins were also upregulated. Though foremost associated with ribosomal protein translation, many EIF3 subunits are also present within the nucleus [29]. Indeed, EIF3E is implicated in DDR pathways, independently of it’s function in protein translation [50] and many other EIF3 proteins are known to be associated with chromatin [29], yet the functional significance of this remains unclear.

## Conclusions

The present study highlights a prominent role for DNA damage in DLB pathology. We provide evidence supporting the occurrence of DNA damage, specifically DSBs within DLB cortical tissue, and demonstrate that in mice the induction of DSBs is a direct consequence of the expression of dysfunctional mutant A30P aSyn. Furthermore, we observed the cytoplasmic accumulation of damaged genomic DNA within LBs. The data suggests a reciprocal relationship, whereby aSyn pathology induces DNA damage, which in turn generate ectopic cytoplasmic genomic material, capable of facilitating cytoplasmic aSyn aggregation. Cellular pathology occurred alongside marked upregulation of DDR, UPS and EIF3 complex proteins. Observed changes in nuclear proteome are supportive of disease induced genomic instability. The overexpression and potentially overactivation of selective DDR components, each have capacity to perturb repair, inducing additional damage and trigger related cell death. Despite the progress in delineating the occurrence of DNA damage and DDR alteration in DLB, it remains to be determined where within these pathways, nuclear aSyn may interact and the specific functional significance of the selective changes in DDR components.

## Supplemental tables

**Supplementary table 1.** Individual details for human tissue cohort. Each case used is listed with an arbitrary case number (Case no) and sex, age (years), post-mortem delay (PMD, hrs), Braak stage, Thal phase, Consortium to Establish a Registry for Alzheimer’s Disease (CERAD), the National Institute of Ageing – Alzheimer’s Association (NIA-AA) criteria, Lewy body (LB) Braak stage and McKeith criteria detailed. Additionally, tissue use for histology A (Hisa-XRCC1+γH2A.X), histology B (Histb-γH2A.X+pS129), Western blot (West), Mass spectrometry (MS) is also provided. NA = not available.

| Case nr | Diagnosis | age | sex | PMI | NFT Braak | Thal Phase | CERAD | NIA-AA | LB Braak | McKeith | Usage |
| --- | --- | --- | --- | --- | --- | --- | --- | --- | --- | --- | --- |
| 1 | Con | 88 | F | 22 | III | 0 | 0 | Not | 0 | No | Hisa+b+West |
| 2 | Con | 60 | M | 60 | 0 | 1 | 0 | Low | 0 | No | Hisa+b+MS+West |
| 3 | Con | 88 | M | 24 | II | 3 | 0 | Low | 0 | No | Hisa+b+MS+West |
| 4 | Con | 85 | M | 45 | III | 4 | 1 | Inter | 0 | No | Hisa+b+MS+West |
| 5 | Con | 90 | F | 63 | II | 0 | 0 | Not | 0 | No | Hisa+b+West |
| 6 | Con | 73 | M | 25 | 0 | 0 | 0 | Not | 0 | No | Hisa+b+MS+West |
| 7 | Con | 72 | M | 60 | II | 5 | 0 | Low | 0 | No | Hisa+b+MS+West |
| 8 | Con | 95 | F | 66 | III | 0 | 0 | Not | 0 | No | Hisa+b+West |
| 9 | Con | 80 | F | 25 | II | 1 | 0 | Low | 0 | No | Hisa+b +MS+West |
| 10 | Con | 94 | F | 15 | II | 1 | 0 | Low | 0 | No | Hisa+b |
| 11 | Con | 81 | M | 43 | II | 0 | 0 | Not | 0 | No | Hisa+b+West |
| 12 | Con | 66 | M | 56 | 0 | 2 | 0 | Low | 0 | No | Hisa+b+MS+West |
| 13 | Con | 99 | F | 5 | II | 0 | 0 | Not | 0 | No | Hisa+MS+West |
| 14 | Con | 74 | F | 74 | III | 0 | 0 | Not | 0 | No | MS+West |
| 15 | DLB | 86 | M | 96 | III | 5 | 2 | Inter | 4 | Limbic | Hisa+b |
| 16 | DLB | 91 | F | 10 | III | 5 | 2 | Inter | 6 | Neo | Hisa+b |
| 17 | DLB | 77 | M | 24 | III | 2 | 0 | Inter | 6 | Neo | Hisa+b+MS+West |
| 18 | DLB | 76 | M | 13 | II | 0 | 0 | Low | NA | Neo | Hisa+b+MS+West |
| 19 | DLB | 72 | M | 89 | III | 0 | 0 | Not | 6 | Neo | Hisa+b+West |
| 20 | DLB | 77 | M | 8 | II | 0 | 0 | Not | NA | Neo | Hisa+b+MS+West |
| 21 | DLB | 81 | M | 21 | III | 4 | 1 | Inter | 6 | Neo | Hisa+b+West |
| 22 | DLB | 75 | M | 18 | II | NA | NA | NA | NA | Limbic | Hisa+b+MS+West |
| 23 | DLB | 81 | M | 26 | III | 3 | 2 | Inter | 6 | Neo | Hisa+b +MS+West |
| 24 | DLB | 71 | M | 68 | II | NA | NA | NA | NA | Neo | Hisa+b+MS+West |
| 25 | DLB | 73 | M | 47 | III | 1 | 0 | Low | 6 | Neo | Hisa+b+West |
| 26 | DLB | 82 | M | 46 | II | 3 | 0 | Low | 6 | Neo | Hisa+b+MS+West |
| 27 | DLB | 78 | M | 8 | III | 4 | 2 | Inter | 6 | Neo | MS+West |
| 28 | DLB | 80 | M | 34 | III | 4 | 1 | Low | 6 | Neo | MS+West |

## Supplemental figures

**Supplementary Figure 1.**
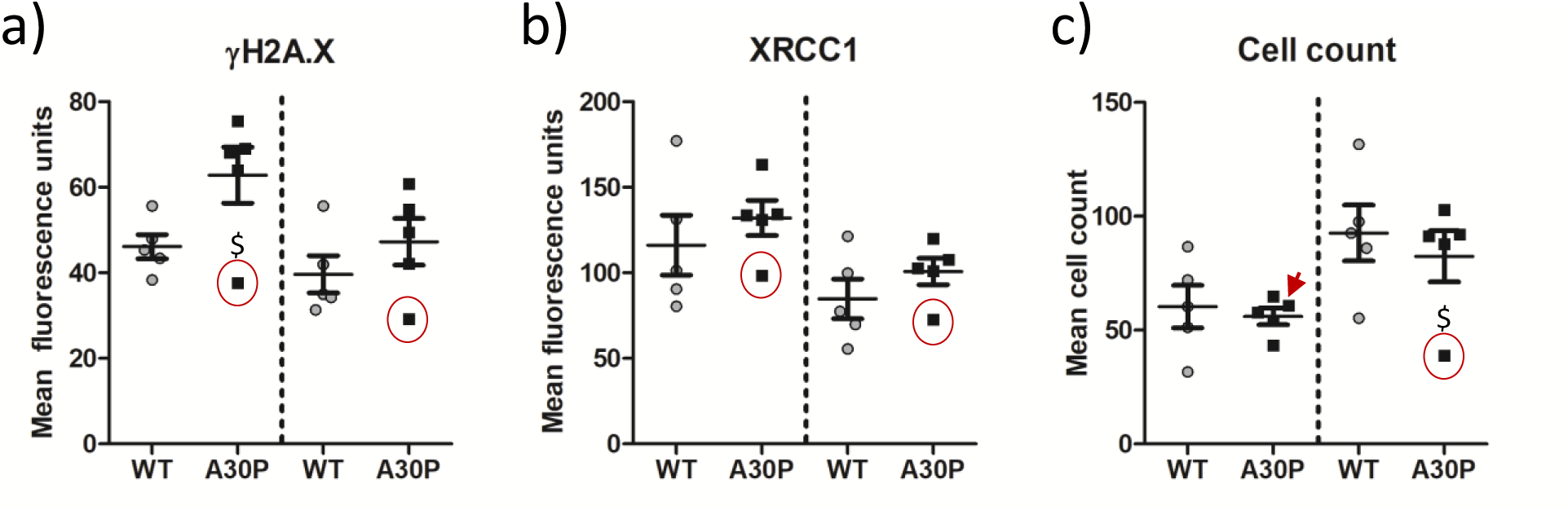
WT and A30P mice full data set including outlier. Identification of statistical outlier with A30P mice was established via Grubbs test as highlighted by red circled data point or arrow. Data is shown for (a). Quantification of neuronal (NeuN +) and non-neuronal (NeuN -) mean γH2A.X (a) XRCC1 (b) and (C) cell count of NeuN+ and NeuN - nuclei signal in WT (n=5) and A30P mice (n=5). Data are expressed in scatterplots with mean±SEM (in b, c and f), $ =p<0.05 in Grubbs outlier test.

**Supplementary Figure 2.**
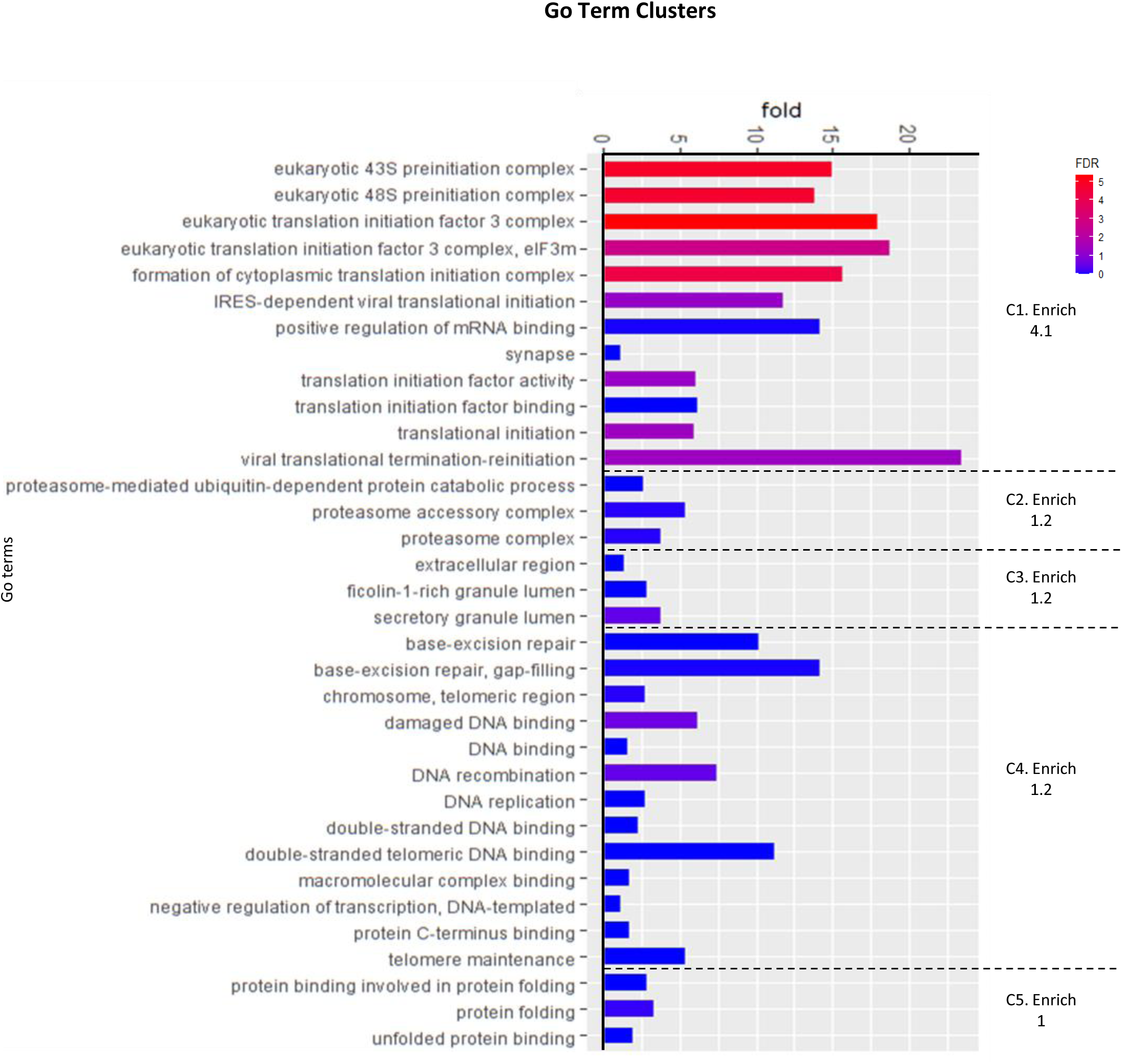
Go term cluster enrichment of the 83 proteins established as elevated in nuclear proteosome of DLB compared to controls. Fold enrichment and significance of individual terms are shown as per -log false discovery rate (FDR). Keywords are grouped into clusters as reported via the DAVID bioinformatic database. Dotted lines denote the boundaries of clusters, with cluster enrichment (C.Enrich) from background reference database shown within each cluster.

## Acknowledgements

Tissue for this study was provided by the Newcastle Brain Tissue Resource which is funded in part by a grant from the UK Medical Research Council (G0400074), by NIHR Newcastle Biomedical Research Centre awarded to the Newcastle upon Tyne NHS Foundation Trust and Newcastle University, and as part of the Brains for Dementia Research Programme jointly funded by Alzheimer’s Research UK and Alzheimer’s Society.

